# KIF16B mediates anterograde transport and modulates lysosomal degradation of the HIV-1 envelope glycoprotein

**DOI:** 10.1101/2023.03.01.530732

**Authors:** Nicholas Weaver, Jason Hammonds, Lingmei Ding, Grigoriy Lerner, Krista Dienger-Stambaugh, Paul Spearman

## Abstract

The human immunodeficiency virus type 1 (HIV-1) envelope glycoprotein (Env) is incorporated into virions at the site of particle assembly on the plasma membrane (PM). The route taken by Env to reach the site of assembly and particle incorporation remains incompletely understood. Following initial delivery to the PM through the secretory pathway, Env is rapidly endocytosed, suggesting that recycling is required for particle incorporation. Endosomes marked by the small GTPase Rab14 have been previously shown to play a role in Env trafficking. Here we examined the role of KIF16B, the molecular motor protein that directs outward movement of Rab14-dependent cargo, in Env trafficking. Env colocalized extensively with KIF16B+ endosomes at the cellular periphery, while expression of a motor-deficient mutant of KIF16B redistributed Env to a perinuclear location. The half-life of Env labeled at the cell surface was markedly reduced in the absence of KIF16B, while a normal half-life was restored through inhibition of lysosomal degradation. In the absence of KIF16B, Env expression on the surface of cells was reduced, leading to a reduction in Env incorporation into particles and a corresponding reduction in particle infectivity. HIV-1 replication in KIF16B knockout cells was substantially reduced as compared to wildtype cells. These results indicate that KIF16B regulates an outward sorting step involved in Env trafficking, thereby limiting lysosomal degradation and enhancing particle incorporation.

**IMPORTANCE:** The HIV-1 envelope glycoprotein is an essential component of HIV-1 particles. The cellular pathways that contribute to incorporation of envelope into particles are not fully understood. Here we identify KIF16B, a motor protein that directs movement from internal compartments toward the plasma membrane, as a host factor that prevents envelope degradation and enhances particle incorporation. This is the first host motor protein identified that contributes to HIV-1 envelope incorporation and replication.

## INTRODUCTION

The HIV-1 Env protein plays critical roles in the viral lifecycle, mediating receptor and coreceptor binding, fusion, and entry into target cells. Env is the principal neutralizing determinant on the surface of virions, and has been a primary focus of vaccine design efforts since the onset of the HIV pandemic. While many aspects of Env structure and function are now understood, the determinants of particle incorporation of Env are not fully defined. HIV-1 Env is synthesized on endoplasmic reticulum (ER)-bound ribosomes as a precursor gp160 protein. Trimerization and initial glycosylation of Env take place in the ER, followed by additional glycan modifications and cleavage of gp160 into gp120 (SU) and gp41 (TM) subunits in the Golgi apparatus. After delivery to the PM, the heterotrimeric Env complex is rapidly endocytosed into early endosomes (1–3). The trafficking events that occur following Env endocytosis are poorly defined, and are the subject of this report.

Clues to Env trafficking and particle incorporation have come from studies of the Env cytoplasmic tail (CT). Long CTs are a characteristic feature of most lentiviral Env proteins. HIV-1, HIV-2, and simian immunodeficiency virus (SIV) have Env CTs of approximately 150 amino acids. The CT of HIV-1 Env contains multiple functional motifs that have been implicated in intracellular trafficking events [reviewed in (4, 5)], suggesting that the CT interacts with host trafficking molecules to direct intracellular trafficking and particle incorporation of Env. In support of the critical role of the CT, Murakami and Freed showed that there are remarkable cell type-specific differences in incorporation of Env with a large deletion in the CT (6). While some cell lines such as MT-4 and 293T cells readily incorporate Env bearing a truncated CT, incorporation into virions in most T cell lines, in primary T lymphocytes, and in monocyte-derived macrophages (MDMs) requires an intact CT (6–9). HeLa cells demonstrate an intermediate phenotype, with a reduction but not complete loss of Env incorporation with a CT truncation (6). The cell type-dependent incorporation of Env with a truncated CT, together with evidence supporting trafficking motifs within the CT, suggests that specific host factors are required for incorporation of Env into particles.

We previously demonstrated that a Rab-related adaptor protein Rab11-FIP1C (FIP1C) plays a role in Env incorporation, and outlined a key role for recycling pathways in this process (10–12). The specific Rab GTPase involved in FIP1C-mediated recycling of Env was shown to be Rab14 (10). Additional evidence supporting a role for Rab14 comes from Hoffman and colleagues, who demonstrated that Env traffics to Rab14+ late endosomes and lysosomes (13). Recycling of Env to the PM was demonstrated in this study, and a potential role for Rab14 in regulating lysosomal degradation vs. delivery to the PM was suggested. Given the accumulating evidence for Rab14 in the intracellular trafficking of Env, we sought to evaluate the role of the molecular motor protein that associates with Rab14, KIF16B.

KIF16B is a member of the kinesin-3 family of molecular motor proteins. KIF16B is a plus end-directed molecular motor that regulates the steady-state distribution of Rab5+ early endosomes and plays a role in recycling of endocytosed cargo to the cell surface (14). A seminal study describing KIF16B function demonstrated that disruption of KIF16B led to redistribution of epidermal growth factor receptor (EGFR) and enhanced its degradation in the lysosome (14). KIF16B has also been implicated in anterograde transport of transferrin receptor and fibroblast growth factor receptor (FGFR), in contributing to MHC class I cross-presentation, and in directing apical transcytosis of cargo in epithelial cells (14–17). KIF16B binds directly to Rab14, and a Rab14-KIF16B complex has been shown to be essential for transport of FGFR and in mediating antigen cross-presentation by dendritic cells (15, 16). Given the role of Rab14 in HIV-1 Env trafficking, we hypothesized that KIF16B-mediated transport plays a role in anterograde trafficking of Env. We found that KIF16B directs Env to early endosomes located at the periphery of cells, while expression of a motor-deficient KIF16B molecule retained Env in perinuclear endosomal compartments. Measurement of the half-life of Env in KIF16B knockout (KO) cells revealed an enhanced rate of Env degradation that could be reversed using an inhibitor of lysosomal acidification. Knockout of KIF16B expression in a T cell line diminished cell surface levels of Env incorporation, Env incorporation into particles, particle infectivity, and viral replication in a spreading infection. These studies implicate KIF16B in anterograde movement of Env away from degradation in the lysosome, allowing more efficient incorporation into HIV-1 particles.

## RESULTS

### Env localizes to KIF16B-positive compartments and is redistributed by dominant-negative KIF16B overexpression

To determine if HIV-1 Env is found in KIF16B-positive endosomal compartments, we co-expressed the HIV-1 NL4-3 Env with KIF16B tagged with yellow fluorescent protein (YFP) in HeLa cells. KIF16B+ compartments were readily observed at the periphery of cells and demonstrated a striking colocalization with Env (Fig. 1A). Expression of KIF16B bearing a mutation in the nucleotide binding motif that disrupts its motor function, KIF16B-S109A-YFP, has been shown to act in a dominant-negative fashion to impair EGFR trafficking (14). Similar to what was seen with EGFR, HIV-1 Env was redistributed by expression of KIF16B-S109A- YFP to a perinuclear location (Fig. 1B). Expression of Env from intact NL4-3 proviral DNA together with wildtype KIF16B led to a peripheral distribution of Env, similar to that seen upon expression of Env alone (Fig. 1C). Expression of intact provirus with KIF16B-S109A- YFP also revealed relocalization of Env to a perinuclear location (Fig. 1D). To quantify the observed difference in subcellular localization, we used a mask 30% the distance from the outer edge at the periphery of the cell to the stained nucleus, and measured Env signal within this mask as compared to total Env signal in the cell. 40 cells were examined for this analysis. The significant differences in peripheral localization shown in the representative images were confirmed quantitatively, as shown in Fig. 1E. Expression of either WT or S109A KIF16B resulted in a high degree of colocalization with Env, as measured by Pearson’s coefficient (Fig. 1F). We noted a somewhat higher degree of colocalization with KIF16B-S109A-YFP expression, which was statistically significant when Env alone was expressed (Fig. 1F). These results largely mirror results reported for KIF16B relocalization of EGFR (14), and support the hypothesis that KIF16B promotes anterograde transport of Env in early endosomal compartments.

**FIG 1.**
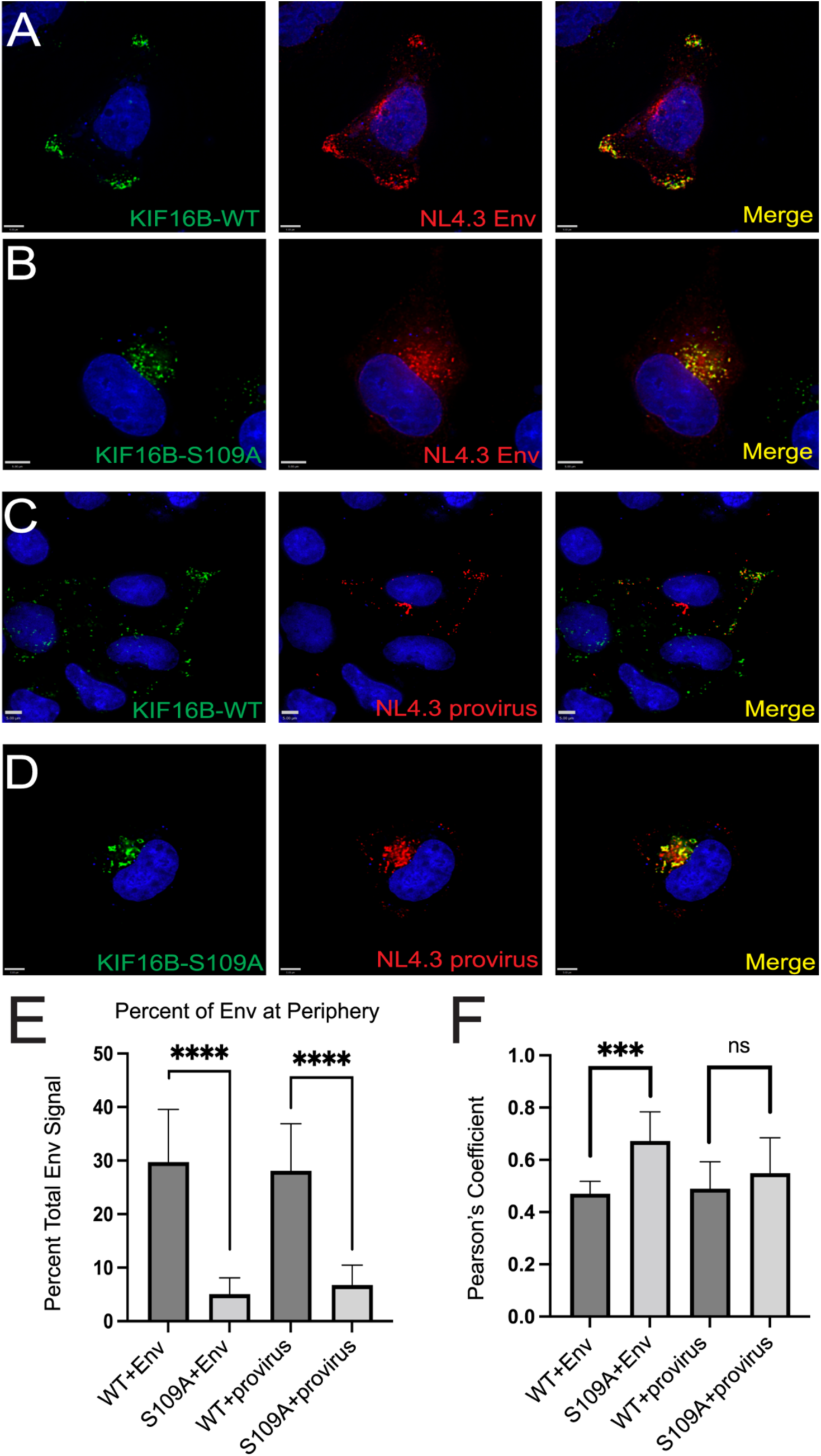
Env localizes to KIF16B-positive compartments and is redistributed by dominant- negative KIF16B. (A) Subcellular distribution of NL4-3 Env when coexpressed with KIF16B-WT-YFP. Cells were fixed and immunolabeled with human neutralizing antibody 2G12 to stain HIV-1 gp120. Green, KIF16B; red, Env; rightmost image, overlay. (B) Distribution of NL4-3 Env when coexpressed with dominant KIF16B-S109A-YFP. Staining is as described for panel A. (C and D) Coexpression of proviral NL4-3 with WT and dominant-negative KIF16B, respectively. Staining for Env as described from A, B. (E) Percent of Env signal at periphery of cells observed for each combination of coexpression shown in A-D. Signal at the periphery was the sum all Env signal within 30% of the distance from edge of cell to the nucleus. Results are shown as means ± SD from a total of 10 representative images. ns = not significant; ***, P < 0.0008; ****, P <0.0001. P values were calculated using the Student’s unpaired t test using GraphPad Prism 9. (F) The degree of colocalization between KIF16B constructs and Env was measured by Pearson’s correlation coefficient equation using Volocity 6.5.1 software after thresholding. Bars represent 5 μm.

### KIF16B fails to redistribute heterologous viral glycoproteins

We considered the possibility that overexpression of KIF16B could divert early endosomes to the periphery of cells in a non-specific manner that would be seen with other viral envelope glycoproteins. To address this possibility, we examined the effect of expression of WT vs. S109A KIF16B on the distribution of three unrelated viral glycoproteins: influenza glycoprotein hemagglutinin (HA), Ebola virus glycoprotein (EBOV GP), and severe acute respiratory syndrome coronavirus 2 spike (SARS-CoV-2 S) protein. As noted previously, WT KIF16B was consistently found at the periphery of cells, while KIF16B-S109A localized near the cell nucleus. No redistribution of HA, EBOV GP, or SARS-CoV-2 S was seen for KIF16B vs. KIF16B-S109A expression (Fig. 2A-F). Furthermore, in contrast to HIV-1 Env, these heterologous viral glycoproteins failed to colocalize significantly with KIF16B as indicated by visual inspection and by determination of a slightly negative Pearson’s Coefficient (supplemental Fig. S1C). These results demonstrate that HIV-1 Env colocalization with and redistribution by KIF16B is not widely generalizable to viral glycoproteins, at least as indicated by examination of a paramyxovirus, filovirus, and coronavirus envelope glycoprotein. Next, we examined the effects of expression of KIF1B, a closely related kinesin mediating plus end-directed trafficking, or its dominant negative (motor-deficient) form KIF1B-Q98L (18), on HIV-1 Env distribution. KIF1B was found at the cellular periphery, while and KIF1B-Q98L localized to perinuclear locations in a manner similar to that of KIF16B and KIF16B-S109A (supplemental Fig. S1A, 1B). However, distribution of HIV-1 Env did not appear to shift to the periphery or perinuclear region upon expression of KIF1B and KIF1B-Q98L, respectively (Fig. S1A, 1B). In contrast to results with KIF16B, Env was not significantly colocalized with either KIF1B protein (Fig. S1C). Together, these results suggest that KIF16B plays a specific role in anterograde trafficking of HIV-1 Env.

**FIG 2.**
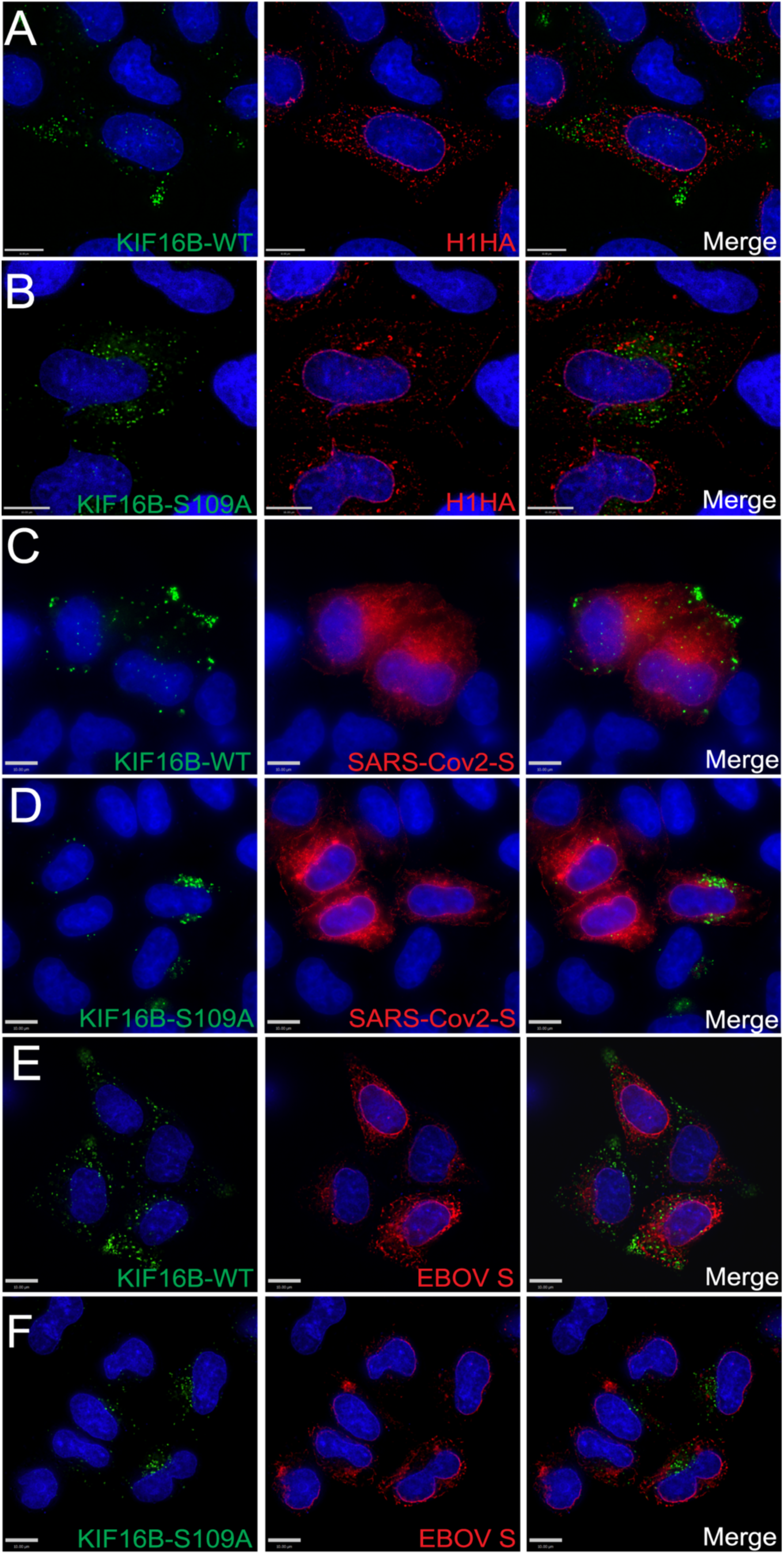
KIF16B and Env colocalization is not generalized to other viral envelope glycoproteins. (A, B) Influenza HA glycoprotein was coexpressed with KIF16B-WT-YFP and KIF16B-S109A-YFP, respectively. Cells were fixed and immunolabeled with 11692- T54 antibody to stain HA. Green, KIF16B; red, HA; rightmost image, overlay. (C, D) SARS-Cov2-S protein was coexpressed with WT and dominant negative KIF16B, respectively. Cells were fixed and immunolabeled with 1A9 to stain spike. Green, KIF16B; red, SARS-Cov2-S; rightmost image, overlay. (E, F) Coexpression of KIF16B-WT-YFP or KIF16B-S109A-YFP with Ebola GP. Cells were fixed and immunostained with 4F3 antibody specific to EBOV GP. Green, KIF16B; red, EBOV GP; rightmost image, overlay. Bars represent 10 μm.

### HIV-1 Env on the PM is endocytosed into KIF16B positive endosomes

The studies described above examined the total pool of Env in the cell, and did not study the endocytosed pool of Env in isolation. In order to determine if Env is delivered into KIF16B positive endosomes directly from the PM, we utilized a novel approach using flourogen activating protein (FAP) tagging (19). The FAP tag was placed within the V2 loop of the ectodomain of JR-FL Env as illustrated in Fig. 3A, creating the FAP-Env construct. To validate this system for use in assessing Env incorporation, we examined particle incorporation of FAP-Env. FAP-Env was co-expressed with Env-deficient provirus, and was efficiently incorporated into virions as indicated by Western blot analysis (supplemental Fig. S2). Combining expression of this construct (FAP-Env) with a flourogen impermeant to the PM (MG-11p, αRED-np1 from Spectragenetics) allowed visualization of the fate of pulse-labeled Env present on the surface of cells. We first co-expressed FAP-Env with KIF16B WT in HeLa cells and fixed the cells after pulse labeling them on ice to visualize the surface staining in the absence of endocytosis. Fig. 3B demonstrates that the cell membrane-impermeable reagent fluoresced upon binding to Env on the cell surface and did not penetrate the PM. We next allowed internalization of FAP-Env for 15 minutes at 37°C after pulse-labeling, and found that surface-labelled FAP-Env on the surface had accumulated into KIF16B-positive compartments (Fig. 3B). We also performed the same experiment with KIF16B S109A and found similar results (Fig. 3C). In order to follow the movement of this FAP-Env population expressed on the cell surface in real-time, we pulse-labeled FAP-Env and recorded live-cell video of Env movement into HeLa cells expressing KIF16B or KIF16B-S109A. Pulse-labeled Env was endocytosed within a few minutes and demonstrated increasing colocalization with KIF16B+ endosomes at the periphery of cells, or with KIF16B-S109A+ endosomes in the perinuclear region of the cell (Fig. 3D and 3E, and supplemental video 1 and 2). These results demonstrate that HIV-1 Env on the surface of the cell is rapidly endocytosed and thereafter accumulates in KIF16B+ endosomes.

**FIG 3.**
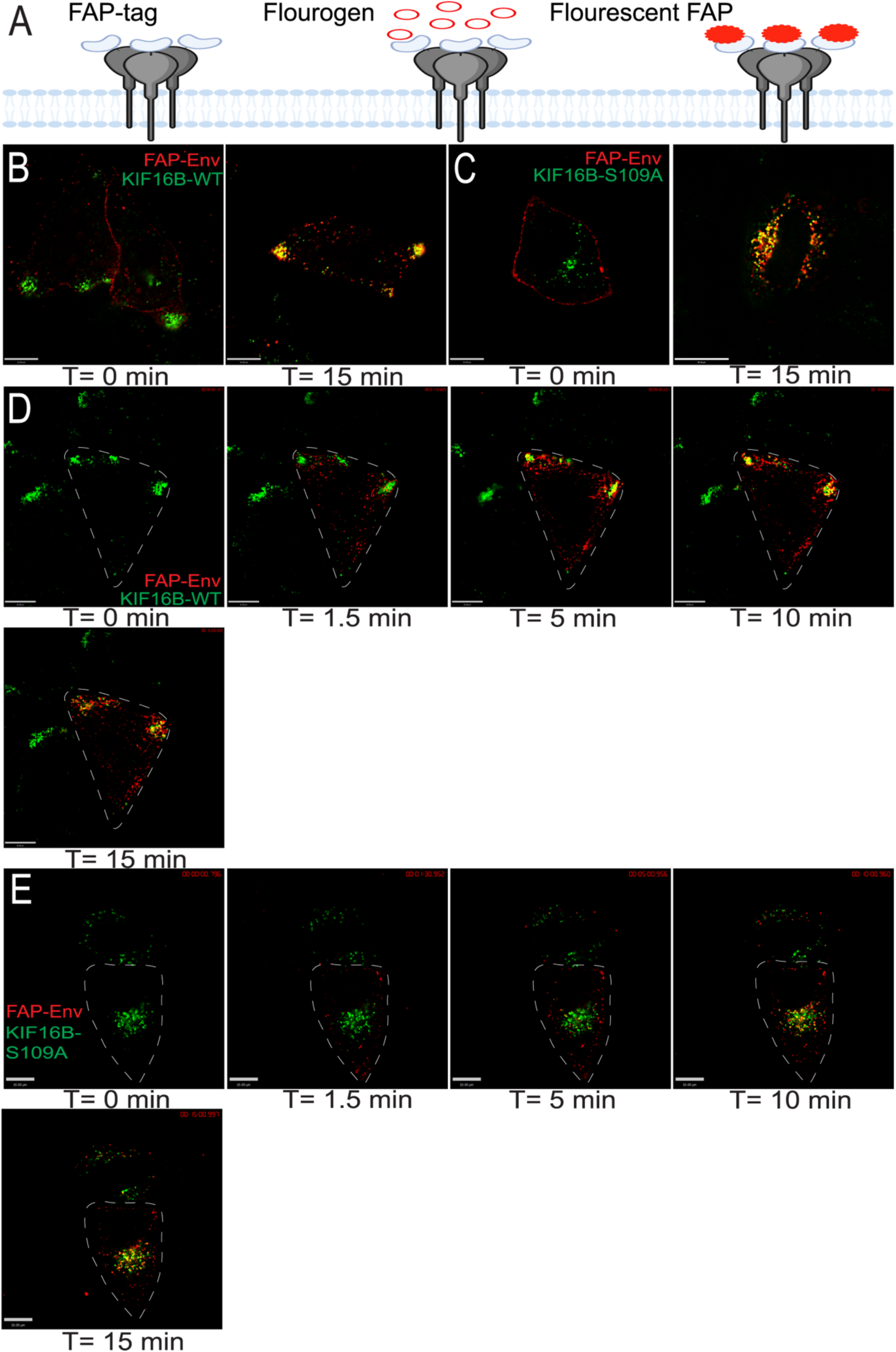
Env on the plasma membrane is endocytosed into KIF16B positive compartments. (A) FAP-Env construct shown as cartoon diagram. The FAP-tag was placed on the V2 loop of the Env ectodomain. This tag is inherently non-fluorescent on the far left. In the middle, the addition of a non-permeable dye is shown. On the far right, the FAP-tag becomes fluorescent, allowing visualization of a pulsed population of surface Env. (B) FAP-Env coexpressed with KIF16B- WT-YFP in HeLa cells. Left image, FAP-Env is pulse labeled for 10 min on ice. Cells were then fixed and imaged immediately to show population of Env on plasma membrane surface. Right image, cells were pulsed as before and then were returned to 37°C incubator for 15 min to allow internalization, then fixed and imaged to show FAP-Env endocytosed into KIF16B-positive compartments. (C) FAP-Env construct co-expressed with KIF16B-S109A-YFP in HeLa cells. Left image is FAP-Env collecting on surface as before. Right image is cell shown after 15 min of internalization as before. (D, E) Frames from live cell imaging video of FAP-Env coexpressed with KIF16B-WT-YFP and KIF16B-S109A-YFP, respectively. Time T= 0 min shows cell before pulse. Subsequent time points show FAP-Env collecting in KIF16B- positive compartments. Dotted lines are a visual aid to represent periphery of cell body. Bars represent 10 μm.

### Deletion of KIF16B leads to enhanced degradation of Env and accumulation within the lysosome

It has been established that KIF16B-mediated transport can divert cellular cargo from default pathways that lead to lysosomal degradation, while depletion of KIF16B enhances their degradation (14, 16) We therefore hypothesized that a loss of normal anterograde trafficking of Env mediated by KIF16B could result in enhanced Env degradation. To evaluate this possibility, we utilized CRISPR/Cas9 technology to knockout KIF16B in both HeLa cells and H9 T cells, followed by characterization of isolated cell clones. Fig. S3 shows characterization of one HeLa and one H9 clone used for subsequent studies, and is consistent with the findings from several clones examined for each cell type. Note the complete absence of KIF16B by Western blotting in supplemental Fig. S3A. We then utilized the KIF16B KO HeLa cell clones together with pulse-labeling of Env using the FAP tag technique described above. FAP-Env was expressed in WT or KIF16B KO cells, and the kinetics of disappearance of total cellular FAP-Env signal was measured over time. These experiments were controlled for the effects of photobleaching on FAP-Env signal as outlined in Materials and Methods. Remarkably, the FAP-Env signal degraded at a considerably faster rate in KIF16B KO cells than in WT cells (images in Fig. 4A vs. Fig. 4B and supplemental video 3). The half-life of endocytosed Env measured in WT HeLa cells by this method was 159.5 minutes (95% C.I. 145.4 – 175.6), while in the absence of KIF16B Env was more rapidly degraded, with a half-life of 56.9 minutes (95% C.I. 54.1 – 59.8). Furthermore, when we incubated KIF16B KO cells with lysosomal inhibitor bafilomyacin-A1 (Baf-A1), the degradation rate in KIF16B KO cells returned to wild-type levels (Fig. 4 A, B, C, E and supplemental video 3). To corroborate the observation that KIF16B knockout leads to enhanced degradation of Env as measured by degradation of FAP-tagged Env, we performed surface pulse-labeling with NHS-biotin in WT and KIF16B KO HeLa cells on ice, followed by quenching of remaining biotin and pulldown of internalized biotinylated proteins with streptavidin beads at multiple timepoints following incubation at 37°C. Detection of internalized Env was then performed using quantitative Western blotting. Figure 4D shows Western blot examples representative of results from five individual experiments for each experimental arm. As compared with WT cells, pulse-labeled Env in KIF16B KO cells was more rapidly degraded (Fig. 4D, compare WT to KO blots). The addition of Baf-A1 reduced the degradation to essentially WT levels (Fig. 4D, 4F). Using repeated experiments, we generated degradation curves for pulsed Env in WT, KO, and KO + Baf-A1 cells that were similar, but not identical, to the curves derived from FAP-Env labeling (Fig. 4F, compare to 4E). Env half-life measured by this method was 154.1 minutes for WT cells, 32.1 minutes for KO cells, and 94.9 minutes for Baf-A1-treated KO cells. While we noted that the results were not identical between the two methods employed, they are in good general agreement, and establish that KIF16B KO leads to more rapid Env degradation that can be rescued by Baf-A1. Together, these results clearly show that KIF16B limits lysosomal degradation of Env, likely through promoting anterograde trafficking.

**FIG 4.**
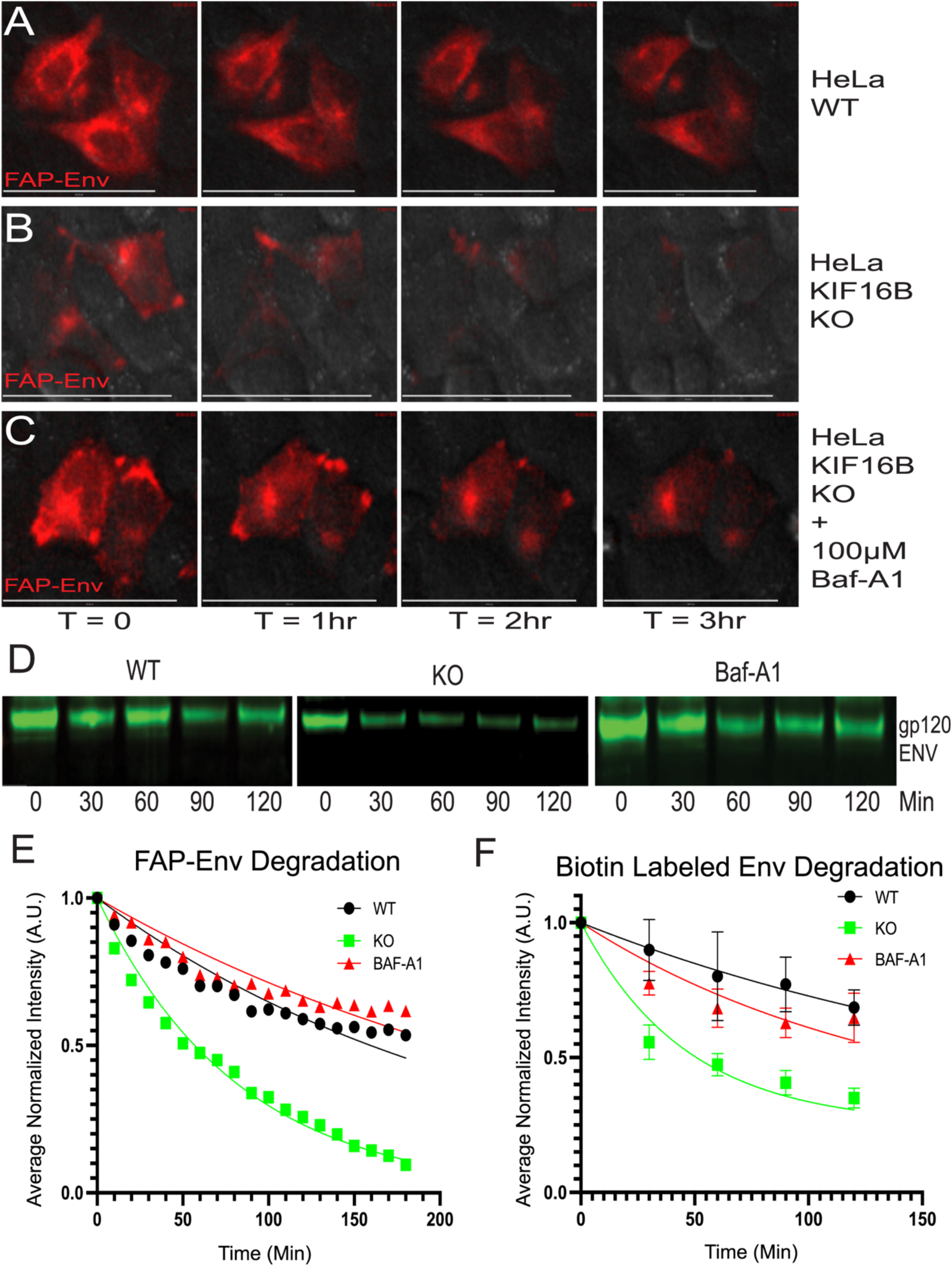
KIF16B KO leads to enhanced degradation of endocytosed Env in the lysosome. (A, B, C) Frames from live cell movies for Fap-Env expressed in (A) WT HeLa cells with vehicle DMSO; (B) KIF16B KO cells with DMSO; or (C) KIF16B KO cells with 100μM Baf- A1. Representative frames are shown for each hour over the time course. (B) Western blots shown are biotinylated Env from surface-labeled cells, with lysates chased over the time course, pulled down with streptavidin beads, and blotted for Env. Labels indicate WT, KIF16B KO, and KIF16B KO + Baf-A1 HeLa cells. (C) Quantification of signal intensity of each time point over the FAP-Env imaging experiment normalized to first frame. (D) Quantification of the Western blot signal for each time point for biotin- streptavidin pulse-chase experiment. Error bars represent SD from 5 independent experiments. Fit curves for both C and D were generated using the Prism 9 non-linear regression one-phase decay algorithm. Bars represent 60 μm.

Given results above with the lysosomal inhibitor Baf-A1, we sought to determine if the absence of KIF16B results in a shift in Env distribution into the lysosomal compartment. Results of immunofluorescence staining were striking. WT HeLa cells revealed punctate lysosomes with a moderate degree of Env colocalization, while intracellular Env was clearly distinguished outside of the lysosome (marked by LAMP1 staining) (Fig. 5A). In KIF16B KO cells, Env colocalization with LAMP1 was dramatically higher (Fig. 5B and 5D). Baf-A1 treatment of KIF16B KO cells did not show significant differences in colocalization of Env and LAMP1 as compared with untreated KO cells, as both showed very high levels of colocalization as quantified by Pearson’s coefficient (Fig. 5C and 5D). Taken together, these results from imaging and pulse-chase analysis suggest that deletion of KIF16B increases the rate of Env degradation due to loss of anterograde movement normally conferred by KIF16B, resulting in shunting to the lysosome for degradation.

**FIG 5.**
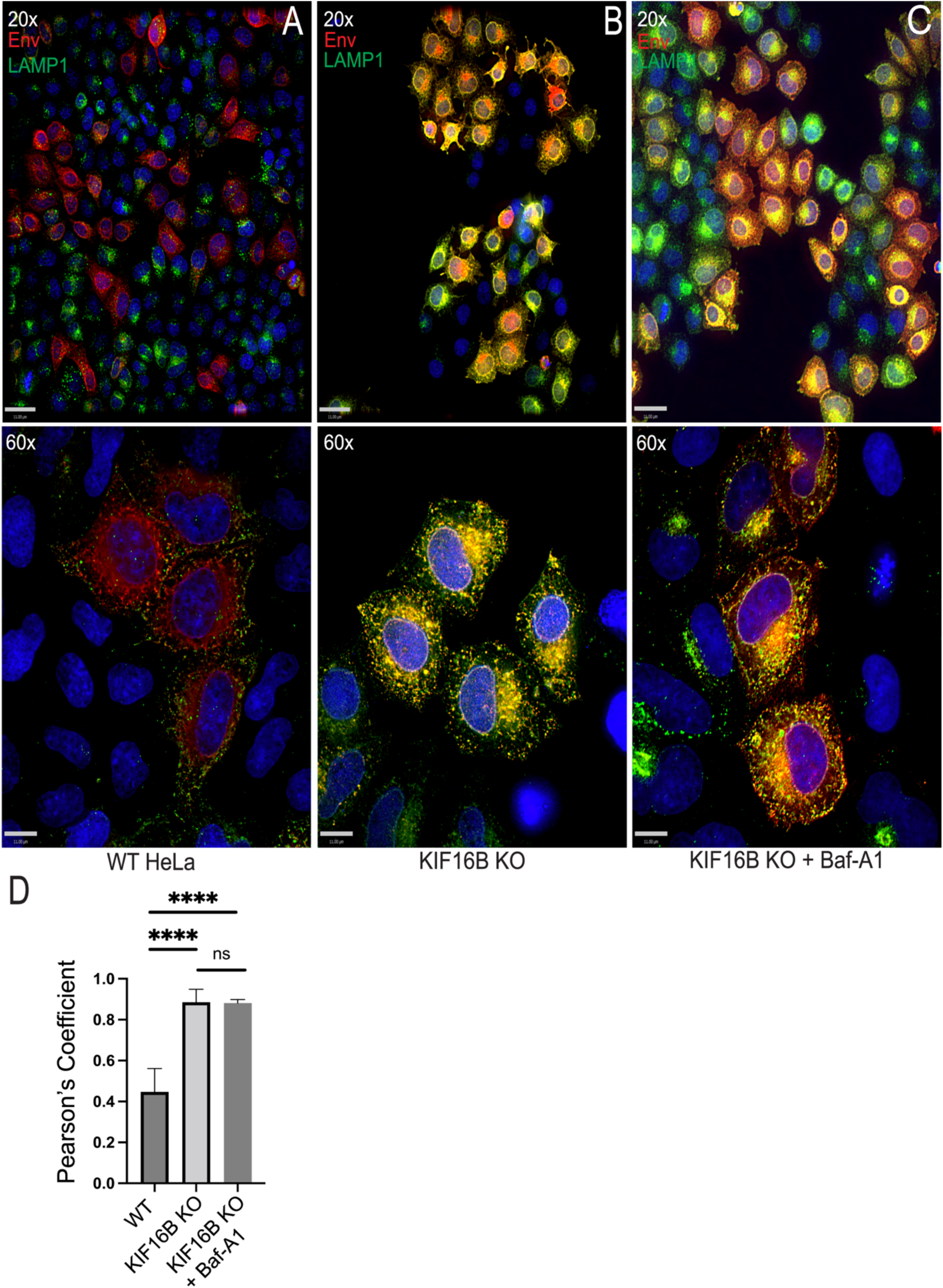
KIF16B KO shunts Env to lysosomal compartments. (A) HIV-1 NL4-3 Env expressed in WT HeLa cells. Top image 20x magnification, bottom image 60x. (B) NL4.3 expressed in KIF16B KO HeLa cells. Top image 20x magnification, bottom image 60x. (C) NL4-3 expressed in KIF16B KO HeLa cells with Baf-A1 added for 3 hours prior to imaging. For all imaging, cells were fixed and immunolabeled for Env with 2G12 antibody and for endogenous LAMP1 with H4A3 antibody. (D) Pearson’s correlation coefficient showing the degree of colocalization between Env and LAMP1 positive compartments between WT, KIF16B KO cells, and WT cells with Baf-A1 in A, B, and C, respectively. Pearson’s correlation coefficient equation using Volocity 6.5.1 software after manual thresholding. Unpaired t-test performed on Prism 9 to get significance values for difference in colocalization. ****, P < 0.0001; *, P = 0.0155. Bars represent 11μm.

### KIF16B deletion leads to defects in Env incorporation, particle infectivity, and viral replication

We next examined the effect of KIF16B depletion on Env trafficking and incorporation into particles. We first assessed surface levels of Env by flow cytometry after infecting WT or KIF16B KO H9 cells with NL4-3 virus. Surface expression levels of Env were consistently diminished in KIF16B KO cells compared with WT cells (Fig. 6A and 6B), consistent with diminished anterograde movement of Env. Total cellular Env measured by flow cytometry was also diminished, but to a somewhat lesser degree than cell surface Env (Fig. 6B). This result is consistent with the enhanced lysosomal degradation of endocytosed Env shown in prior results. Viruses were then purified from supernatants of infected WT and KIF16B KO H9 cells, in order to examine levels of Env incorporation into particles. Viral particle Env content was significantly reduced in KIF16B KO cells compared to WT H9 (>50% reduction from wild-type levels as quantified by LI- COR Image Studio Software, n = 5) (Fig. 6C). The leftmost blot in Fig. 6C shows particle Env content normalized by equal p24 content loading of the lanes. When cell lysates were examined following multiple rounds of replication in WT or KIF16B KO cells and normalized for total protein, there was a reduction of all viral proteins, reflecting diminished replication (Fig. 6C, cell lysate). Notably, however, production of gp160 in KIF16B KO cells was not diminished, as indicated by Western blot of lysates from infected cells examined after a single round of replication (Fig. 6C, rightmost blot). This suggests that the reduced levels of Env seen on the PM and on particles derived from KIF16B KO cells were not due to an Env production defect. To evaluate the significance of the diminished particle incorporation of Env induced by KIF16B KO, we next examined particle infectivity and viral replication. Consistent with the observed reduction of Env on particles, the infectivity of particles released from KIF16B KO cells was consistently reduced (Fig. 6D). We noted that the reduction in Env incorporation shown in Fig. 6C and the reduction in particle infectivity in 6D were consistent but are incomplete, suggesting that although KIF16B alters the level of Env incorporation, it is not absolutely required for incorporation of Env into particles. To determine the significance of this level of Env reduction on viral replication, we infected WT H9 and KIF16B KO cells and measured viral replication in a spreading infection. Infection of KIF16B KO H9 cells resulted in a significant delay in replication and a reduced peak of released virus as compared with WT H9 cells (Fig. 6E). We conclude from these results that the anterograde movement of Env by KIF16B plays a role in Env incorporation and viral replication. Loss of KIF16B results in lower levels of cell surface Env and a corresponding loss of Env incorporation into particles.

**FIG 6.**
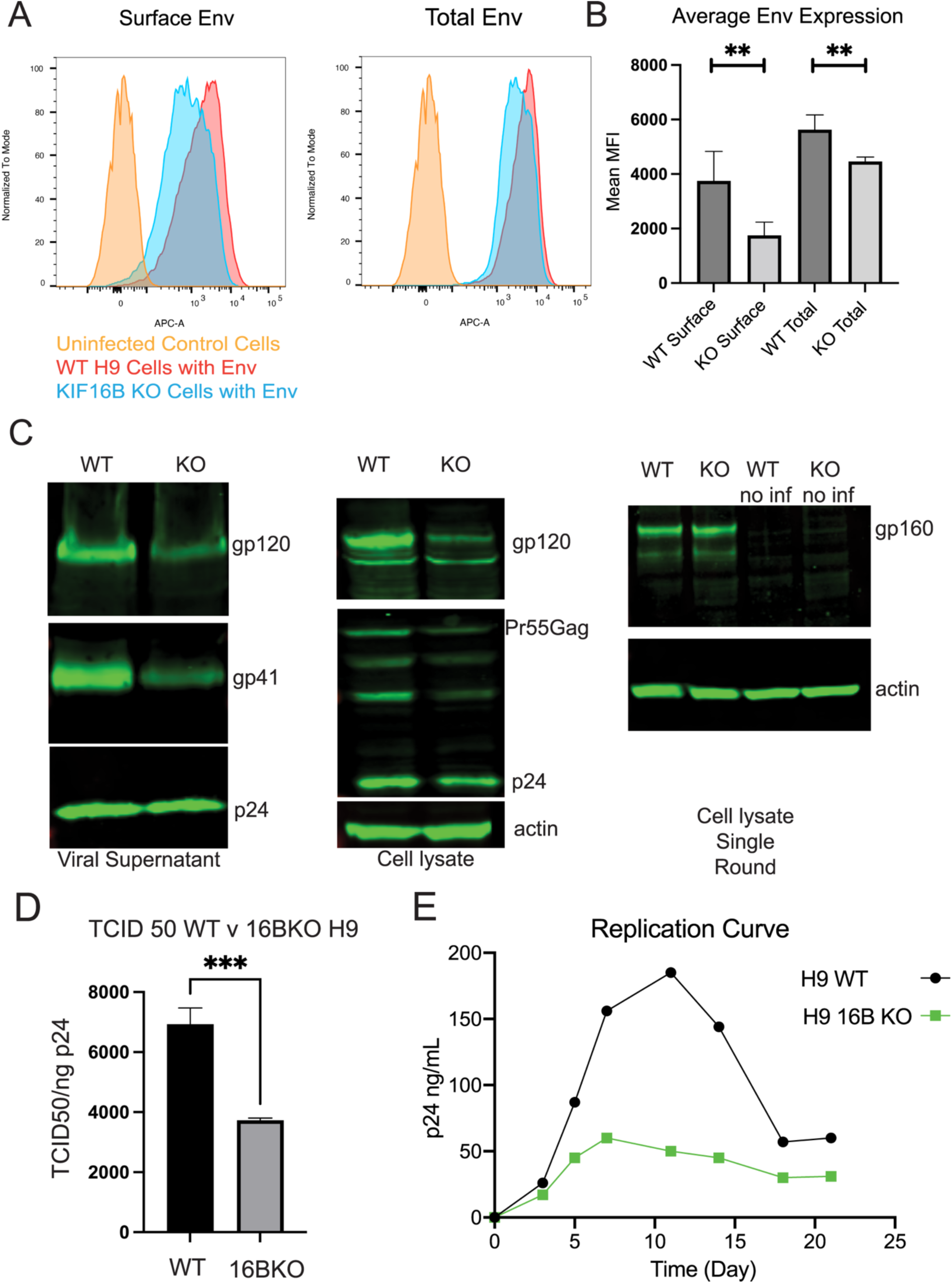
KIF16B depletion leads to reduction of Env on the cell surface, particle incorporation, infectivity, and viral replication. (A) Flow plots for cell surface staining (non-permeabilized) and total Env content (permeabilized) in WT H9 and KIF16B KO H9 cells. Cells were fixed and stained for Env using anti-gp120 2G12 antibody. (B) Quantification of flow cytometry mean fluorescent intensity of Env expression on surface and total cell averaged across 5 independent experiments. Unpaired t-test performed on Prism 9 to get significance values between surface vs total. **, P < 0.005 (C) WT and KIF16B H9 cells were infected with VSV-G-pseudotyped NL4-3. Shown are Western blots of Env and Gag content for particles in viral supernatant (left) and cell lysates (middle) collected on day 5 following infection. Note that loading was normalized for p24 content (supernatants) or for total cellular protein (cell lysates). Rightmost blot, single-round infection of WT and KIF16B KO H9 cells infected with VSV-G-pseudotyped NL4-3 at an MOI of 0.5 and harvested for cell lysate analysis after 24 hours. Uninfected WT and KO cell lysates were run on the same blot as controls. Blotting was performed with 2G12 antibody against HIV- 1 gp160 and ACTN05 (Invitrogen) against actin as a loading control. (D) Infectivity of viral particles released from WT or KIF16B KO H9 cells from 3 independent experiments evaluated using TZM-bl indicator cells. (E) WT (black line) and KIF16B KO (green line) were infected with VSV-G-pseudotyped NL4-3 virus. Viral growth was monitored over time by detection of p24 in the culture supernatant.

## DISCUSSION

Molecular motor proteins are essential to the normal function of eukaryotic cells. The cytoplasm of cells is dense, crowded with proteins, vesicles, nucleotides, and organelles. Free diffusion of large molecules within the cytoplasm is restricted, and cannot be relied upon to move these cellular materials with the efficiency and accuracy required for proper cell function. Instead, the cell must use specialized molecular motors. These motors – kinesins, dyneins, and myosins – use adenosine triphosphate (ATP) to power movement of cargo along cytoskeletal structures. Kinesins and dyneins utilize microtubule networks as their pathways for transporting cellular cargo. With some exceptions, kinesins move in an anterograde direction along microtubules. Dyneins, in contrast, move in a retrograde direction toward the minus end of microtubules often localized at the microtubule-organizing center (MTOC). Kinesins and dyneins are most commonly structured with an N-terminal motor head that is responsible for powering processive, step-like movement of motor subunits, attached via a middle stalk region to a C-terminal tail composed of light and heavy chains that bind the cellular cargo (20–23) .

Viruses hijack host cellular pathways in order to perform essential steps in their lifecycle, including co-opting molecular motor-driven intracellular transport (24–27). HIV-1 has been shown to utilize motor proteins at several steps of replication (27, 28). Following entry to a cell, the conical core of HIV-1 has been shown to bind to dynein and to use the KIF5 motor protein to successfully traffic towards the nucleus (29, 30). Depletion of the KIF5B motor prevents nuclear entry and infection of HIV-1 in a CA-dependent manner (31). In studies of outward trafficking of Gag, KIF4 has been shown to directly interact with Gag in both yeast two-hybrid and co-immunoprecipitation assays (32, 33). Additionally, disruption of KIF4 slowed progression of Gag through trafficking intermediates, and depletion of KIF4 increased degradation of Gag that was reversed upon reintroduction of KIF4 (34). However, no motor protein has previously been shown to interact with the HIV-1 Env protein or participate in its trafficking within host cells. Because Env that reaches the cell surface through the secretory pathway is rapidly endocytosed and must subsequently traffic in an anterograde manner to return to the PM for particle incorporation, we hypothesized in this study that HIV-1 would also likely use a kinesin or kinesins for outward trafficking of Env. Several pieces of evidence implicate Rab14 as an important outward trafficking factor for Env. Our group has shown that Rab14 mediates FIP1C-dependent recycling of Env and plays a role in Env incorporation (10). Hoffman and colleagues similarly found that Env colocalizes to Rab14+ endosomal compartments in T cell lines, and unexpectedly found that these compartments shared some characteristics with late endosomes and lysosomes (13). KIF16B is an effector of Rab14-mediated trafficking that binds directly to Rab14 (15). In this study, we demonstrate that KIF16B plays a role in Env trafficking and in the ability of HIV-1 to form fully infectious particles. Similar to other reported cargo of KIF16B such as EGFR, Env colocalized strongly with KIF16B at the periphery of cells. Motor mutant dominant negative KIF16B redistributed Env to a perinuclear position, entirely consistent with what had been seen for EGFR. Video microscopy of pulse-labeled Env expressed on the surface of cells in this study indicated that Env on the PM is endocytosed into KIF16B- positive compartments. In the absence of KIF16B, Env was shifted to the lysosome, and the half-life of Env that had been endocytosed from the PM was significantly reduced. Taken together, these studies strongly suggest that KIF16B mediates anterograde movement of Env and limits the pool of Env that is shunted to the lysosome for degradation. To our knowledge, this is the first evidence for a motor protein contributing to Env trafficking and particle incorporation.

Data from this study support the model shown in Fig. 7. According to this model, Env is transported through the secretory pathway to the PM and subsequently endocytosed to the early or recycling compartments, following which Env can be directed in an anterograde fashion by KIF16B or can follow a default pathway leading to fusion with the lysosome (Fig. 7A). In the absence of KIF16B, a larger pool of endocytosed Env is shunted to the lysosome, as indicated by the relative size of the arrows in Fig. 7B. The fraction of the endocytosed pool of Env that is directed to the periphery and is subsequently available for incorporation into particles is diminished in the absence of KIF16B, resulting in diminished particle incorporation, particle infectivity, and reduced replication. Fig. 7 for simplicity does not include other potential trafficking routes of Env, such as retrograde trafficking to the Golgi and the potential for alternative routes for Env to recycle to the PM.

**FIG 7.**
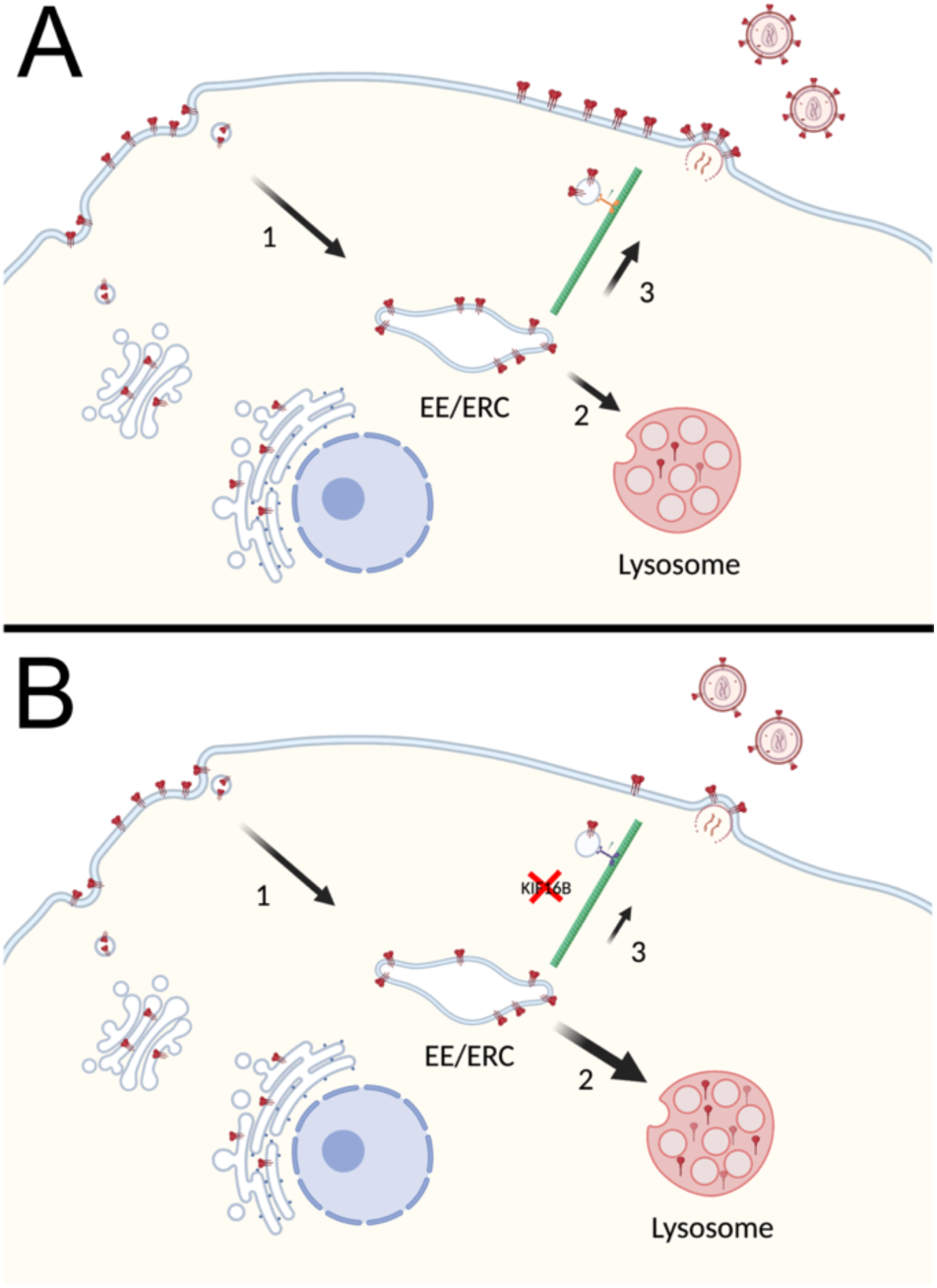
Model of Env Trafficking with (A) and without (B) KIF16B. (A) In a wild-type cell, Env is produced in the rough endoplasmic reticulum and follows the secretory pathway to the cell surface. Env is then endocytosed into early endosomes (path #1 in the figure), which later may sort to the endosomal recycling compartment (ERC) for recycling. This compartment is depicted as “EE/ERC”. A fraction of Env in EEs can progress to fuse with lysosomes, leading to lysosomal degradation in path #2. In path #3, Env is directed in an anterograde manner to the PM, where it is available for incorporation into viral particles. Path #3 is the relevant pathway that is partially dependent upon KIF16B. (B) In a cell with KIF16B KO, a diminished fraction of Env is delivered to the PM from the EE/ERC via path #3. This results in an increase in the amount of Env shunted to the lysosome (path #2, illustrated as broader arrow) and enhanced degradation of Env. The change in the distribution of endocytosed Env is shown graphically as fewer trimers on the budding and mature virion, as well as fewer trimers on the PM and more trimers in the lysosome. Figure created with BioRender.com.

Four models for Env incorporation into HIV particles have been proposed by the Freed laboratory (5). Passive incorporation at the PM is not likely to be the correct model in nonpermissive cell types including primary T cells and macrophages, as Env with a full-length CT is excluded in these cell types. This model remains a potential in permissive cell lines that do not require the CT (4). The identification of host factors such as Rab11- FIP1C, Rab14, and KIF16B that influence particle incorporation favors the Gag-Env co-targeting model, whereby Env is directed to specific microdomains on the PM that are enriched in Gag molecules and serve as sites of particle budding. At present there is no evidence favoring an indirect interaction model with an adaptor or linker connecting Gag to Env. The direct Gag-Env interaction model has been supported by two biochemical studies (35, 36), and by evidence that maturation is required before Env can mediate membrane fusion, while CT truncation relieves the constraint on fusion posed by the immature Gag lattice (9, 37, 38). Trimerization of MA has been shown to be required for Env incorporation, while some mutations in MA that result in Env exclusion from particles can be rescued by compensatory mutations at the trimer interface (39, 40). Furthermore, MA trimerization-impaired mutants can be rescued by substitutions that simultaneously restore Env incorporation (41). This has added substantially to the evidence favoring an interaction of the CT with MA in the form of MA trimers, and supports a model whereby the structural integrity of the MA lattice is an important determinant of Env incorporation (41). Results in the present study showing a role for kinesin-mediated trafficking of Env do not negate the importance of structural integrity of the MA lattice for particle incorporation of Env. Trafficking to the site of the developing Gag lattice mediated by host factor interactions with the CT may be a necessary step preceding a direct Gag-Env interaction. We expect that a defect in anterograde trafficking and increase in Env degradation in the lysosome would diminish the delivery of Env to sites of assembly and reduce the potential interaction of Env with the Gag lattice.

This study provides new information regarding the half-life of Env trimers following endocytosis from the PM. A prior study employing autoradiographic pulse-labeling of total cellular Env indicated that cellular gp160 and gp120 are degraded over a period of hours, and that inhibition of lysosomal acidification prevents gp160 degradation (42). Degradation of gp160 has been shown to not only occur in the lysosome, but also within the biosynthetic pathway within the ER or Golgi compartments (43, 44). However, these studies labeled all Env populations within the cell, whereas the current study utilized two methods of cell surface pulse-labeling followed by chase periods to measure the half-life of Env trimers that had reached the plasma membrane. We found that endocytosed Env has a half-life of 150-160 minutes, and that the absence of KIF16B promotes much more rapid degradation (57 or 32 minutes, measured by FAP-Env degradation or surface biotinylation, respectively). An important role for lysosomal degradation of endocytosed Env is suggested by the reversal of the enhanced rates of degradation in KIF16B KO cells by Baf-A1. A model in which endocytosed Env is shunted to the lysosome for degradation in the absence of KIF16B is also supported by the increased appearance of Env in lysosomal compartments in KIF16B KO cells.

An important caveat for our conclusions is that a complete KIF16B KO resulted in an incomplete reduction of Env incorporation and particle infectivity. Env incorporation and particle infectivity were diminished by approximately 50%. This suggests that kinesins other than KIF16B may also be involved in trafficking Env to the PM, or that there are multiple pathways whereby Env may be recycled to the surface. Indeed, it is known that successful movement of cargo within cells is often accomplished by groups of motors rather than just one (45). Retrograde trafficking of Env from early endosomes to the Golgi may play a role in Env incorporation into particles, as depletion of components of the retromer complex modulated Env distribution and particle incorporation (46). However, the identification of KIF16B as a regulator of Env trafficking and degradation defines another component of the host machinery regulating Env incorporation. Kinesins have been proposed as potential therapeutic targets in some areas of medicine such as cancer and chronic pain (47–49). Although targeting KIF16B function globally would be disruptive to many cell processes, it is possible that a more directed approach disrupting KIF16B- mediated Env trafficking can be developed. In conclusion, we identify KIF16B as an anterograde motor involved in Env trafficking and particle incorporation. Future studies will be needed to fully elucidate the complex pathways taken by Env to reach the site of particle assembly and enable particle incorporation.

## MATERIALS AND METHODS

### Cells and Plasmids

HeLa cells were obtained from the American Type Culture Collection (ATCC; CRM-CCL-2). TZM-bl cells were obtained through the NIH AIDS Reagent Program, Division of AIDS, NIAID, NIH, from John C. Kappes, Xiaoyun Wu, and Tranzyme, Inc. The H9 T cell line was obtained from ATCC (HTB-176), and cells were maintained in RPMI medium 1640 supplemented with 10% fetal bovine serum (FBS), 2 mM Glutamine, and antibiotics. HeLa and TZM-bl cells were maintained in DMEM (Dulbecco’s modified Eagle medium) containing 10% FBS and 2 mM penicillin-streptomycin. pNL4-3 proviral plasmid was obtained through the NIH AIDS Reagent Program, Division of AIDS, NIAID, NIH, from Malcolm Martin. Env expression plasmid pIIINL4Env was a gift kindly provided by Dr. Eric Freed (HIV DRP, NCI, NIH) (6). The Influenza A vaccine strain of A/Brisbane/02/2018 (H1N1) HA sequence was used to construct a human codon-optimized gene which was synthesized and cloned into the BamHI and EcoRI sites of pcDNA3.1+ by GenScript. The SARS-CoV-2 spike gene (Wuhan) was synthesized by Genscript for Dr. Biao He (University of Georgia) and gifted to our lab. The human codon-optimized full length SARS-CoV-2 spike gene was subcloned into the KpnI and BamHI sites of the plasmid pcDNA5TO/Puro. pcDNA5/TO- puro was constructed from backbone pCDNA5/TO (ThermoFisher Scientific) as previously described (50). The EBOV S protein plasmid pcDNA5/TO-puro EBOV GP has been previously described (51). Plasmid pcDNA5/TO JR-FLoptgp160 expresses a codon-optimized JR-FL *env* gene in the pcDNA5/TO vector backbone (Thermo Fisher Scientific, Waltham, MA) as previously described (12). Overlap-extension PCR was used to create novel Xho1 and Pst1 sites within the exposed portion of the V2 loop of JR-FL. The fluorogen-activated peptide (FAP) sequence from pMFAP-alpha2 (Spectragenetics, Pittsburgh, PA) was PCR amplified as an Xho1-Pst1 fragment and ligated into pcDNA5/TO JR-FLoptgp160, placing the FAP peptide in-frame within the V2 loop. A complete cloning schema with primers is available upon request. The WT KIF16B-YFP and dominant KIF16B-S109A-YFP were kindly provided by Sebastian Hoepfner (14). Expression plasmids for Flag-KIF1Bβ and Flag- KIF1Bβ Q98L were kindly provided Susanne Schlisio and described in Fell et. al. 2017 (18).

### Viral infectivity and multi-round replication assays

H9 T cells were infected with VSV G-pseudotyped NL4-3 virus at an MOI of 0.2. Cells and supernatants were harvested 4-5 days after infection for analysis by Western blotting, p24 antigen quantitation, and infectivity assays. Infectivity of cell culture supernatants was measured using TZM-bl indicator cells following p24 normalization as previously described (52). Replications assays were performed by taking aliquots of the supernatant after infection every 3 days for 21 days. Supernatant was then used in a p24 ELISA assay to graph the amount of virus produced over the course of viral growth and replication. Murine anti-p24 capture antibody 183-H12-5C (CA183) was obtained from Bruce Chesebro and Kathy Wehrly through the NIH AIDS Research and Reference Reagent Program, and the p24 ELISA assay was performed as previously described (53).

### Western blotting, antibodies, and immunostaining

For analysis of viral protein content, viruses were pelleted from cell culture supernatants through a 20% sucrose cushion at 35,000 × g for 3 h at 4°C; cells were harvested and pelleted at 400 × g. Virus pellets and cell lysates were dissolved in radioimmunoprecipitation assay (RIPA) buffer (Thermo Scientific) containing protease inhibitors. Cell lysates for blotting KIF16B were prepared with a hypotonic lysis buffer consisting of 10mM Hepes, 1.5mM MgCL_2_, 10 mM KCl in PBS on ice for 15 minutes, vortexed with 10% IGEPAL CA630 (Sigma) to break open cells, and then centrifuged to pellet out nuclei. Virus pellets and cell lysates were separated on 4 to 12% polyacrylamide gels, transferred to nitrocellulose, and subjected to Western blotting using antibodies outlined below. Goat polyclonal antibody AHP2204, from Bio-Rad AbD Serotec (Oxford, UK)(D7324), was used for detecting HIV-1 gp120 for Western blotting. Recombinant human monoclonal antibody 2F5 from Polymun scientific (Klosterneuburg, Austria) was used to detect HIV-1 gp41 in Western blots. Actin was detected with Invitrogen Actin antibody ACTN05. KIF16B antibody used was polycolonal mouse from Abnova (H00055614-B01P). Mouse anti-p24 monoclonal CA-183 (provided by Bruce Chesebro and Kathy Wehrly through the NIH AIDS Research and Reference Reagent Program) was used to detect p24 and Pr55^Gag^ in Western blotting and for enzyme-linked immunosorbent assay (ELISA) to measure HIV-1 Gag in virus-containing culture supernatant and virus pellets; the capture ELISA was performed as previously described (54). IRDye goat anti-mouse, IRDye rabbit anti-goat, and IRDye goat anti-rabbit secondary antibodies from LiCor Biosciences (Lincoln, NE) were used for Western blotting. All blots were acquired and analyzed using the LiCor Odyssey infrared detection system. The following antibodies were utilized in immunofluorescence assays: human anti-gp120 antibody IgG1 2G12 from Polymun scientific (A002), monoclonal ANTI-FLAG® antibody produced in rabbit from Sigma-Aldrich (F2555), rabbit polyclonal anti- HA1 antibody from Sino Biological (Beijing, China) (11692-T54), mouse monoclonal SARS-CoV / SARS-CoV-2 (COVID-19) spike antibody from Genetex (Irvine, CA) (1A9), mouse anti- EBOV GP monoclonal antibody 4F3 from IBT Bioservices (Rockville, MD), anti-LAMP1 mouse monoclonal antibody from BD Biosciences (Franklin Lakes, NJ) (H4A3).

### Env degradation assays

n order to determine degradation rates of Env microscopically using our FAP-tagged Env, we plated 100k HeLa WT or KIF16B KO HeLa cells in wells of Ibidi 4 well μ-Slide plate (Ibidi) and then transfected overnight with the FAP-Env construct. In one of the KO cell wells, 100μM concentration of Bafilomycin A1 (Baf-A1, Invivogen) was employed while the WT and other KO cell wells received an equal amount of DMSO. Three hours later, live cell imaging was conducted following a 10 min pulse label incubation with FAP reagent αRED-np1 (Spectragenetics) in 5% CO_2_ at 37°C chamber on Widefield Nikon Ti2 inverted SpectraX (Cincinnati Children’s Confocal Core). Data from these movies was extracted on Volocity imaging software (Quorum Technologies) to acquire the signal intensity of each ROI drawn around individual cells analyzed in the field of view over the time course. The averaged normalized signal intensity was then calculated using the method outlined in Alber et. al. to attain the degradation curve and half-life of the FAP-tagged Env protein (55). For protein analysis of Env degradation by Western blotting, we plated 1.75 million WT or KIF16B KO HeLa cells in 10cm dishes (5 for WT, 10 for KO). The next morning, 3ug of pIII-Env and 250 ng of psv72-Tat protein were cotransfected into each plate. Following 24 hours of incubation, 100μM Baf-A1 was added to 5 of the KO dishes, while the other KO and WT dishes received an equal amount of DMSO. Three hours later a 1 mM solution of biotin/PBS was applied to the cells and allowed to label surface proteins for 15 min. This labeling reaction was quenched with 1x TBS solution for 5 min. Cells were then washed with PBS and returned to normal media. Cell plates were then put on ice and in a 4°C cold room to stop endocytosis at 0, 30, 60, 90, 120 min after biotin labeling and were then subsequently scrapped and harvested for cell lysates in RIPA buffer. Cell lysates were then IP overnight with 60μL of Pierce™ High Capacity NeutrAvidin™ Agarose (Thermo-Fischer). Beads were then collected in Pierce™ Spin Columns (Thermo-Fischer) and protein was then eluted off the beads at 95°C in 1x Laemli buffer with reducing agent. After 2000 x g spin for 3 min equal amounts of elution were run for each time point on a 4-12% SDS-Page gel and then analyzed by immunoblotting for gp120.

### Image acquisition and analysis

HeLa cells were plated in MatTek 35mm poly–d-lysine-treated dishes (Brooke Knapp MatTek) overnight and then were then transfected with the indicated constructs using jetPRIME (Polyplus) per manufacturer’s instructions. After a 24 h incubation, cells were rinsed with phosphate-buffered saline (PBS) three times and fixed in 4% paraformaldehyde for 10 min at room temperature. Cells were washed with PBS three times with mild agitation after fixation, and then permeabilized for 10 min with 0.2% Triton X-100, followed by washing with PBS. Dako blocking solution was applied to cells for 30 min under gentle shaking conditions. Cells were then incubated with the appropriate primary antibody detailed above diluted in Dako (Agilent) antibody diluent to 1:500. Species appropriate Alexa-flour (Thermo-Fischer) secondary antibody was diluted in Dako antibody diluent to 1:1,000. Images were acquired on a widefield DeltaVision Elite High Resolution microscope (GE Healthcare). Percent of total Env signal at the periphery of cells was calculated using measurements from Volocity 6.5.1 imaging analysis software. Briefly, regions of interest were drawn around the entire cell, around 30% of the distance from the edge of the cell signal to the nucleus (inner 70% of total cell area), and around the nucleus. Peripheral signal was calculated by subtracting the middle two regions from the sum of signal from the total cell region, giving a percentage of peripheral signal as all signal in the last 30% of the cell body towards the edge. Quantitation of colocalization was performed using Volocity software features by derivation of Pearson’s correlation coefficient after manual thresholding.

### Flow cytometry for HIV-1 Env expression on cell surface

For cell surface staining, approximately 5 million H9 cells were infected overnight with 1000 ng p24 of VSV-G-pseudotyped NL4.3. The next day, cells were washed once with PBS and plated in a six well plate in RPMI for 48 hours. Infected cells were stained for viability with zombie violet dye (BioLegend) diluted 1:500 in PBS, followed by fixation with 4% paraformaldehyde. Cells were blocked with Dako protein block then stained for cell surface Env with 2G12 anti-gp120 directly conjugated to APC (Abcam, ab201807) and diluted 1:100 in Dako antibody diluent (10ug/mL final concentration) for 2 hours. 2G12 was washed out twice with PBS followed by permeabilization with 0.2% triton-X 100. Lastly, cells were stained for total infected cells using p24 antibody, KC-57 FITC diluted 1:100 in Dako antibody diluent. Cells were washed twice with PBS then resuspended in MACS (Miltenyi Biotec) buffer and analyzed using the BD FACS Canto II.

### CRISPR KO strategy

CRISPR-Cas9 technology was used to knockout KIF16B in both HeLa and H9 cell lines. Using the Crispor.tefor.net open access website and algorithm designed to find areas of the genome to target with guide RNAs (gRNA), we targeted Exon 13 of KIF16B using sequence from NCBI (Gene ID 55614, Assebmly GCF_000001405.40, Chr 20, location NC_000020.11) and found the stretch of DNA with the highest score to target for deletion being CTCACGGATAAGTTTGACGT. We ordered sense (CACCGCTCACGGATAAGTTTGACGT) and anti-sense (AAACACGTCAAACTTATCCGTGAGC) gRNA oligos + PAM sequence from Integrated DNA Technologies (IDT) to clone into the lentiCRISPR v2 that was a gift from Feng Zhang (Addgene plasmid # 52961 ; http://n2t.net/addgene:52961 ; RRID:Addgene_52961) using the target guide sequence protocol from Sanjana Lab (http://sanjanalab.org/library/protocol_lentiOligo.pdf) (56) After transformation of MAX Efficiency™ Stbl2™ Competent Cells (Thermo-Fischer) and DNA harvest, we packaged the plasmid into a lentivirus using sPAX2 and PMGD2 and transduced our HeLa and H9 cell lines. Puromycin selection and limiting dilution were used to obtain clones knocked out for KIF16B (Fig. S1).

## ACKNOWLEDGMENTS

This work was supported by 5R01AI150486-24

The funders had no role in study design, data collection and interpretation, or the decision to submit the work for publication.

Flow cytometry was performed using the Cincinnati Children’s Hospital Medical Center (CCHMC) Flow Cytometry Core. Live cell imaging of was conducted in the Confocal Imaging Center at CCHMC. DNA sequencing was performed at the DNA Sequencing core at CCHMC.

---

**Fig S1.**
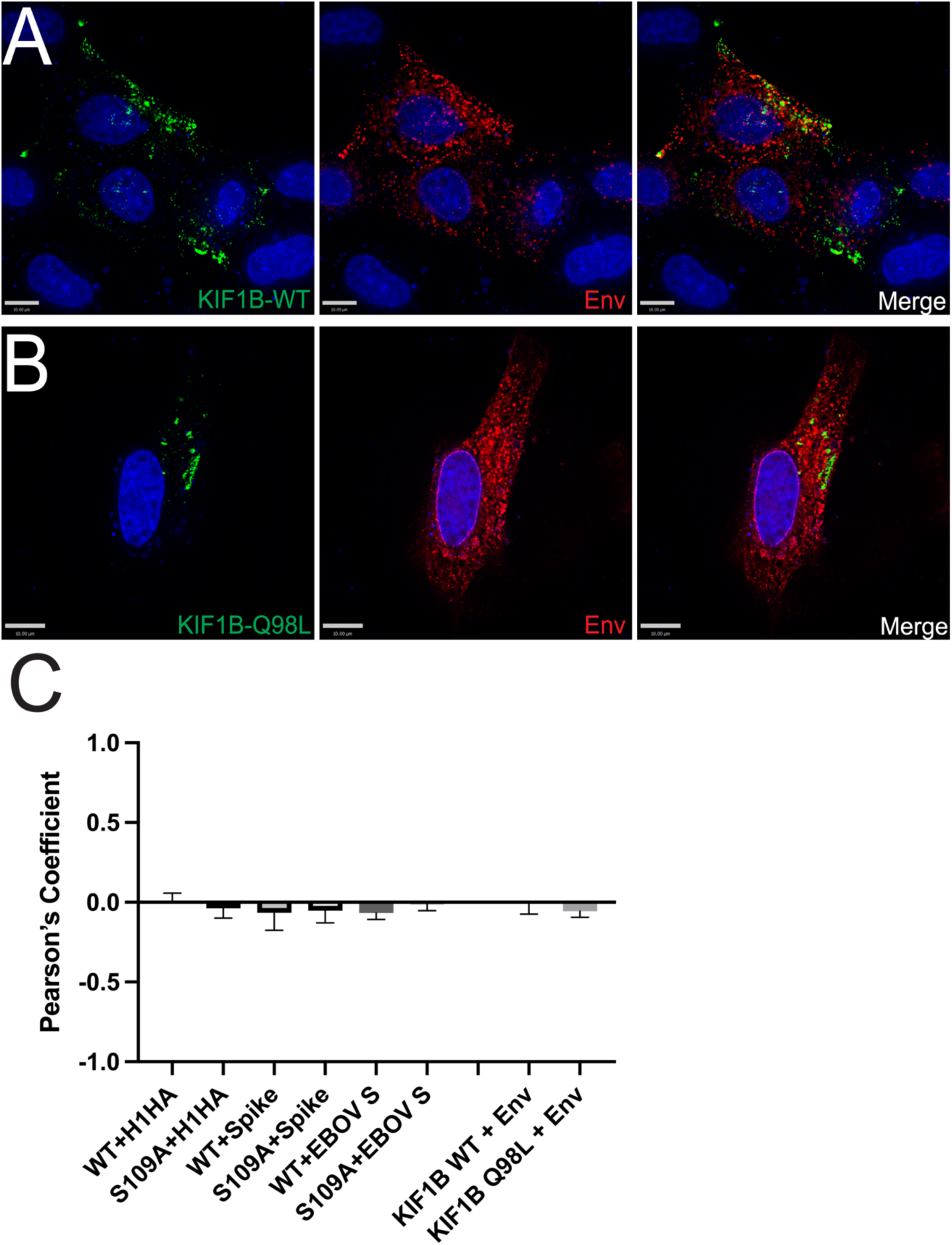
(A, B) Coexpression of KIF1B WT and KIF1B Q98L (dominant-negative) with HIV- 1 Env. Cells were fixed and immunostained with 2G12 antibody specific to the Env glycoprotein. Green, KIF1B; red, Env; rightmost image, overlay. (C) Degree of colocalization for all viral glycoproteins included in Fig 2 with KIF16B (left six bars); and colocalization of Env with KIF1B (rightmost two bars), measured by Pearson’s correlation coefficient equation using Volocity 6.5.1 software after thresholding. Results are shown as means ± SD from a total of 5 representative images. Bars represent 10 μm.

**Fig S2.**
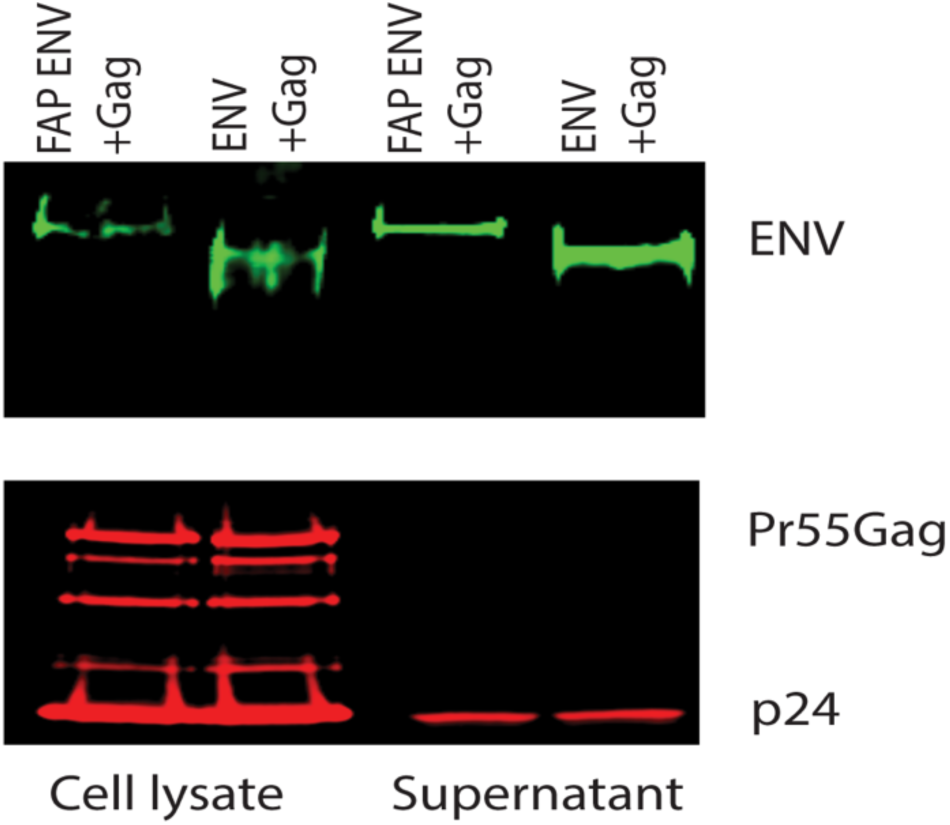
FAP-tagged Env or WT JR-FL Env were co-expressed with Env-deficient provirus. Western blot shows Env in cell lysates and on pelleted particles from supernatants. Env was blotted with 2G12 and Gag detected with CA183.

**Fig S3.**
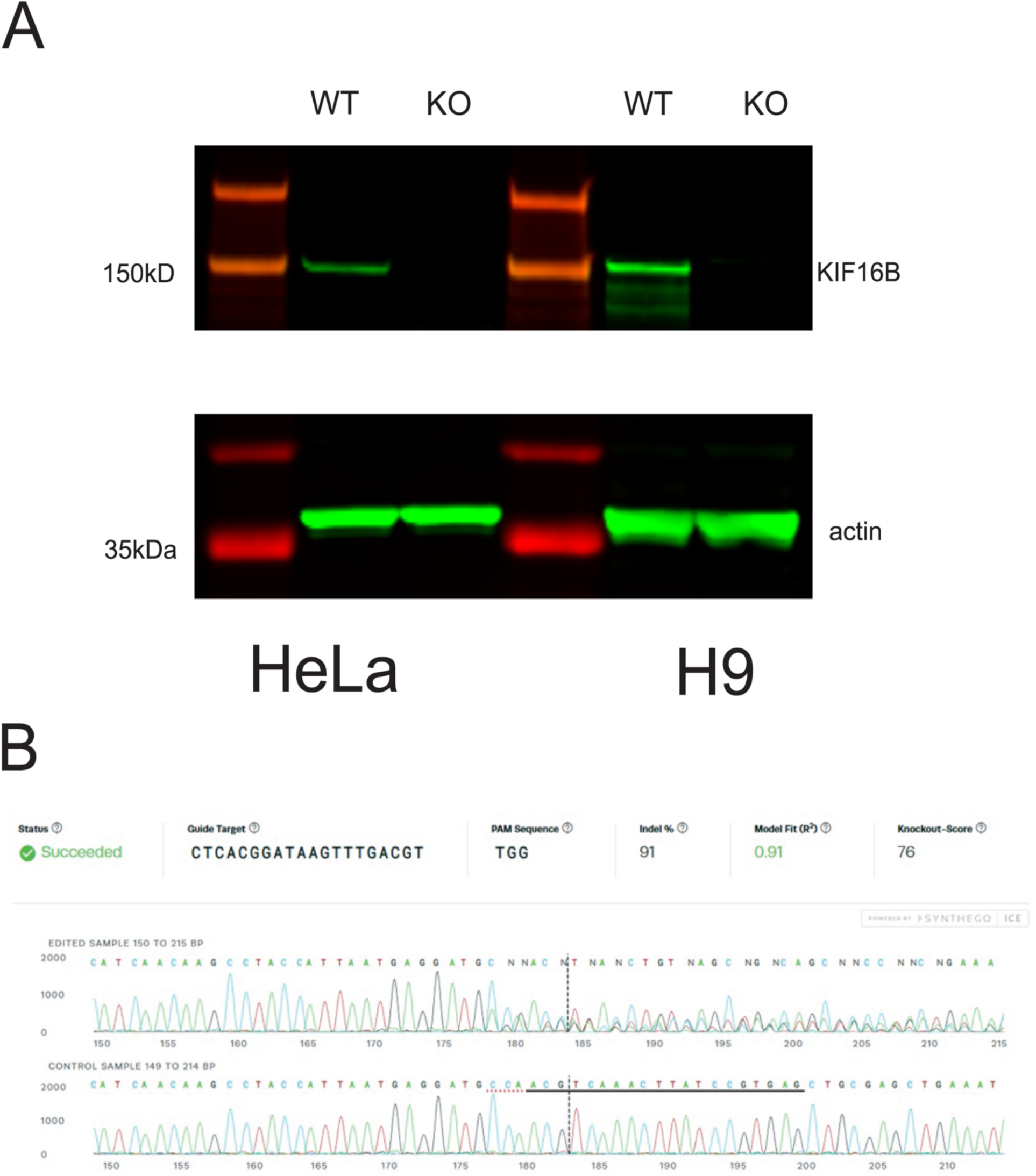
(A) Representative blots of WT and KIF16B KO HeLa (left) and H9 (right) cell lines blotted for KIF16B and actin loading control showing complete protein knock out in both cell lines. (B) Sequence of genomic DNA from HeLa cells showing CRISPR-Cas9 indel. Dotted line indicates cut site of CRISPR-Cas9 complex. Edited sample sequence on top for KIF16B KO cells, the control sample on bottom was the reference sequence from WT cells that were the same cells used to make the knockout clones. Sequence analyzed by ICE analysis via the Synthego software application.

