## Supplemental Video legends for "KIF16B mediates anterograde transport and modulates lysosomal degradation of the HIV-1 envelope glycoprotein"

**Supplemental Video 1** FAP-Env is endocytosed into WT KIF16B positive compartments. FAP-Env construct was co-expressed with WT KIF16B-YFP in HeLa cells. The next day, cells were imaged in a live cell chamber at 37°F with 5% $CO_2$. Impermeable FAP reagent was added to the cell media at 1.5 minutes and is seen upon addition. Over the course of 15 minutes, this surface Env is endocytosed readily into KIF16B positive compartments. Red, FAP-Env. Green, KIF16B. Bar is 10μm.

**Supplemental Video 2** FAP-Env is endocytosed into S109A dominant-negative KIF16B positive compartments. FAP-Env construct was co-expressed with S109A-KIF16B-YFP in HeLa cells. The next day, cells were imaged in a live cell chamber at 37°F with 5% $CO_2$. Impermeable FAP reagent was added to the cell media at 1.5 minutes and is seen upon addition. Over the course of 15 minutes, Env from the surface is endocytosed and transverses the cytoplasm into S109A-KIF16B positive compartments near the nucleus. Red, FAP-Env. Green, KIF16B. Bar is 10μm.

**Supplemental Video 3** KIF16B knockout in HeLa cells enhances FAP-Env degradation, which is recovered by lysosomal inhibitor BAF-A1. (A, B, C) FAP-Env is expressed in (A) WT HeLa cells with vehicle DMSO (B) KIF16B KO HeLa cells with DMSO and (C) KIF16B KO plus 100μM BAF-A1 cells (DMSO or BAF-A1 is placed on cells 3 hours before imaging) overnight prior to imaging. Live cell imaging performed at 37°F with 5% $CO_2$. Impermeable FAP reagent was added to the cell media immediately prior to imaging. Time course of experiment is shown in upper-right as 3 hours. Red, FAP-Env. Bars are 60μm.
